# Four novel Picornaviruses detected in Magellanic Penguins (*Spheniscus magellanicus*) in Chile

**DOI:** 10.1101/2020.10.26.356485

**Authors:** Juliette Hayer, Michelle Wille, Alejandro Font, Marcelo González-Aravena, Helene Norder, Maja Malmberg

**Affiliations:** Department of Animal Breeding and Genetics, Swedish University of Agricultural Sciences, Uppsala, Sweden; Marie Bashir Institute for Infectious Diseases and Biosecurity, School of Life and Environmental Sciences and School of Medical Sciences, The University of Sydney, Sydney, Australia; Instituto Antártico Chileno, Plaza Muñoz Gamero 1055, Punta Arenas, Chile; Department of Infectious Diseases/Virology, Institute of Biomedicine, Sahlgrenska Academy, University of Gothenburg, Sweden; Region Västra Götaland, Sahlgrenska University Hospital, Department of Clinical Microbiology, Gothenburg, Sweden; Department of Biomedical Sciences and Veterinary Public Health, Swedish University of Agricultural Sciences, Uppsala, Sweden

**Keywords:** Sphenisciformes, penguins, Picornaviridae, viral metagenomics, Hepatovirus

## Abstract

Members of the *Picornaviridae* comprise a significant burden on the poultry industry, causing diseases such as gastroenteritis and hepatitis. However, with the advent of metagenomics, a number of picornaviruses have now been revealed in apparently healthy wild birds. In this study, we identified four novel viruses belonging to the family *Picornaviridae* in healthy Magellanic Penguins (*Spheniscus magellanicus*), a near threatened species found along the coastlines of temperate South America. We collected 107 faecal samples from 72 individual penguins. Twelve samples were initially sequenced by high throughout sequencing with metagenomics approach. All samples were subsequently screened by PCR for these new viruses, and approximately 20% of the penguins were infected with at least one of these viruses, and seven individuals were co-infected with two or more. The viruses were distantly related to members of the genera Hepatoviruses, Tremoviruses and unassigned viruses from Antarctic Penguins and Red-Crowned Cranes. Further, they had more than 60% amino acid divergence from other picornaviruses, and therefore likely constitute novel genera. That these four novel viruses were abundant among the sampled penguins, suggests Magellanic Penguins may be a reservoir for several picornaviruses belonging to different genera. Our results demonstrate the vast undersampling of wild birds for viruses, and we expect the discovery of numerous avian viruses that are related to Hepatoviruses and Tremoviruses in the future.

**Importance:** Recent work has demonstrated that Antarctic penguins of the genus *Pygoscelis* are hosts for an array of viral species. However, beyond these Antarctic penguin species, very little is known about the viral diversity or ecology in this highly charismatic avian order. Through metagenomics we identified four novel viruses belonging to the *Picornaviridae* family in faecal samples from Magellanic Penguins. These highly divergent viruses, each possibly representing novel genera, are related to members of the Hepatovirus, Tremovirus genera, and unassigned picornaviruses described from Antarctic Penguin and Red-crowned Cranes. By PCR these novel viruses were shown to be common in Magellanic Penguins, indicating that penguins may play a key role in their epidemiology and evolution. Overall, we encourage further sampling to reveal virus diversity, ecology, and evolution in these unique avian taxa.

## Introduction

Penguins (Order: Sphenisciformes) are unique in the avian world. Through evolution, they have acquired numerous physiological adaptations to exploit and thrive in marine environments. Penguins are found throughout the temperate regions of the Southern Hemisphere, ranging from the Antarctic continent to as far north as the Galapagos Islands. Despite population declines of many penguin species, largely due to climate change and overfishing of their prey, little is known about the pathogens and parasites harboured by these birds. Indeed, infections with various microorganisms are known to play a role in reducing avian populations, such as substantial declines of native Hawaiian birds due to avian malaria, negative survival effects of albatross due to avian cholera, and an negative survival effect of a number of North American passerine species due to infection with West Nile Virus (1–4).

To date, studies of viral presence and prevalence among penguins has been limited, opportunistic, and related to sick birds in nature (eg (5–7)) or in rehabilitation centres (eg (8)). Further, studies of viruses in penguins are highly biased towards the charismatic Antarctic Penguins, with serology studies dating back to the 1970’s (9). Due to technological limitations, *i.e.* both serology and PCR based studies allow for the assessment of only described viruses, very little progress has been made revealing the viral communities in these species. This has changed dramatically with the rise of metagenomics and metatranscriptomics (9, 10), wherein novel and highly divergent viral species may be described. As a result, since 2015, more than 25 different novel viruses have been described in Antarctic and sub-Antarctic penguins (9–11). These viruses include members from the *Adenoviridae, Astroviridae, Caliciviridae, Circoviridae, Coronaviridae, Herpesviridae, Paramyxoviridae, Orthomyxovirdae, Polyomaviridae, Papillomaviridae, Picornaviridae, Picobirnaviridae* and *Reoviridae* (9, 10, 12–15). Beyond Antarctica, evaluation of penguins for viruses is haphazard, although many of the same viral families have been detected through both virology and serology studies of Magellanic Penguins (*Spheniscus magellanicus*), African Penguins (*Spheniscus demersus*), Little Blue Penguins (*Eudyptula minor*), and Galapagos Penguins (*Spheniscus mendiculus*) (6–8, 16–23). It is clear from these studies that the viral diversity of penguins is vastly underappreciated, and thus the role of penguins as potential hosts for an array of viruses are yet to be revealed.

Through metagenomics, a number of novel picornaviruses have recently been described in samples from wild birds. These viruses fall into the genera *Megrivirus, Sapelovirus, Avihepatovirus*, and other highly divergent viruses with unassigned genera (24–29). In penguins, seven picornaviruses have been described: Ross Virus, Scott Virus, Amundsen Virus (Hepatovirus/Tremovirus-like), Shirase Virus (Sinicivirus-like), Wedell Virus (Pigeon Picornavirus B-like), Pingu Virus (Gallivirus-like) and Penguin Megrivirus, all of which have been isolated from apparently healthy Antarctic penguins (10, 13, 30). Until recently, avian picornaviruses were exclusively associated with morbidity and mortality in poultry and other birds (pigeons and passerines) (31, 32). Through an increased effort in investigating the virome in healthy wild birds, we are beginning to rewrite the narrative of many viral families. There is mounting evidence to suggest that picornaviruses are not exclusively disease causing, but rather that many picornavirus species detected in wild birds are not associated with any signs of disease.

In this study we aimed to identify and characterise viruses of Magellanic Penguins. We sampled faeces from Magellanic Penguins breeding on the Magdalena island, an important breeding colony for this species, situated in southern Chile, and we used a combination of high-throughput metagenomic sequencing, followed by PCR screening to disentangle virus diversity and prevalence. The detection of four novel viruses at high prevalence in penguins without obvious disease, provides further evidence that penguins are reservoir hosts for a multitude of viruses belonging to different virus families and genera, including numerous viruses yet to be revealed. Finally, this finding has important evolutionary implications for the emergence/evolution of the picornavirus lineage comprising Hepatovirus and Tremoviruses.

## Results

### Sequence data and assembly

We performed metagenomics sequencing of 12 faecal samples collected from Magellanic Penguins in a colony on Magdalena Island, Chile.

High-throughput sequencing of the sample produced 25,357,939 reads (range 517,072-3,558,413 reads) with a mean length of 229 bp representing a total of 5.86 gigabases of sequence data. After quality control and filtering using PRINSEQ 24,027,073 reads (range 463,974-3,438,3911 reads) remained (5.75 Gb). Of these we assembled an average of 10,228 contigs, of which 4% comprised viral contigs (Table S1). Reads have been deposited to the European Nucleotide Archive (ENA), accession number PRJEB40660.

For all samples, about 25% of the reads could be taxonomically classified. For 7 samples, greater than 50% of the reads could be classified. The majority of these reads were microbial, *i.e.* bacterial or viral; however, a number of penguin (host) reads were also identified. For 2 samples, is_034_015 and is_034_016 viral reads were found in 33.8% and 17.3% of all reads. A vast majority of those were classified as viruses belonging to the *Picornaviridae* family, and three previously not described viral genomes were assembled. From sample is_034_018, 33% (397 out of 1,179 contigs) were classified as viral, of which 313 were classified as *Picornaviridae.*

Overall at the contig level, more than 25% of *de novo* assembled contigs were successfully classified.Eight samples had greater than 50% of the contigs classified as bacterial, viral or chordates.

### Four novel, divergent picornaviruses in Magellanic Penguins

We revealed four novel picornaviruses in faecal samples from Magellanic Penguins: Sphenifaro virus, Sphenigellan virus, Sphenimaju virus, Sphenilena virus. These viruses were recovered from 3 different genomic libraries corresponding to 3 samples from individuals. Two viruses (Spenifaro virus and Sphenilena virus) were identified in a single sample, *i.e.* from the same penguin. Overall, the abundance (proportion of reads based on read mapping back to assemblies) of each virus was low (<1%) with the exception of Sphenimaju virus, which comprised 58.57% of all reads in the sample (is_034_015). The sample is_034_016 that contained two different viruses (Sphenifaro virus and Sphenilena virus) had low abundance for both viruses. Each of these viruses comprised 0.29% and 0.21% of the reads in the sample, respectively (Table 1).

**Table 1:** Metadata for the four novel hepato-like viruses revealed in this study.

| Sample Name | Virus Name | Top hit | Blastn percent identity | Length | Number of reads in sample | Number of reads (proportion of reads in sample) |
| --- | --- | --- | --- | --- | --- | --- |
| is_034_016 | Sphenifaro virus | AUW34301<br>Picornaviridae<br>red crowned crane | 37.91% | 7201 bp | 916,871 | 2,687 (0.29%) |
| is_034_018 | Sphenigellan virus | YP_009164030<br>Phopivirus | 38.76% | 7384 bp | 1,090,489 | 1,238 (0.11%) |
| is_034_015 | Sphenimaju virus | YP_009179216<br>Bat hepatovirus | 38.37% | 7441 bp | 2,689,813 | 1,575,605 (58.57%) |
| is_034_016 | Sphenilena virus | YP_009215780<br>Tupaia hepatovirus A | 35.83% | 7048bp | 916,871 | 1,919 (0.21%) |

These new viruses, recovered from the same penguin colony at the same time, are highly divergent from each other, sharing only 32-43% similarity at the amino acid level, suggesting that they represent not only four different species but potentially four different genera. Three of the viruses were most similar to members in the genus *Hepatovirus* in the *Picornaviridae* family when the genomes were analysed by Blastx. However, at the amino acid level the similarity was low, ranging from 35-37% (Table 1). The fourth virus, Sphenifaro virus, was most similar to an unassigned virus described in Red-crowned Cranes (*Grus japonensis)* when using Blastx, although as with the Blast results of the other viruses, the amino acid percentage identity was low (37.91%). By phylogenetic analysis these viruses were found in the same major lineage as members of the *Hepatovirus* and *Tremovirus* genera, in addition to newly described viruses from Antarctic Penguins (10) and Red-crowned Cranes (29) (Figure 1, Figure S1). However, as with the blast results, the viruses revealed here shared <30% amino acid similarity in the P1 region to other viruses in this lineage. Further, all four viruses had long branch lengths in the phylogeny, all this together suggest that each of these new picornaviruses may represent new genera in the *Picornaviridae* family demonstrating the vast undersampling of viruses in this part of the tree.

**Figure 1:**
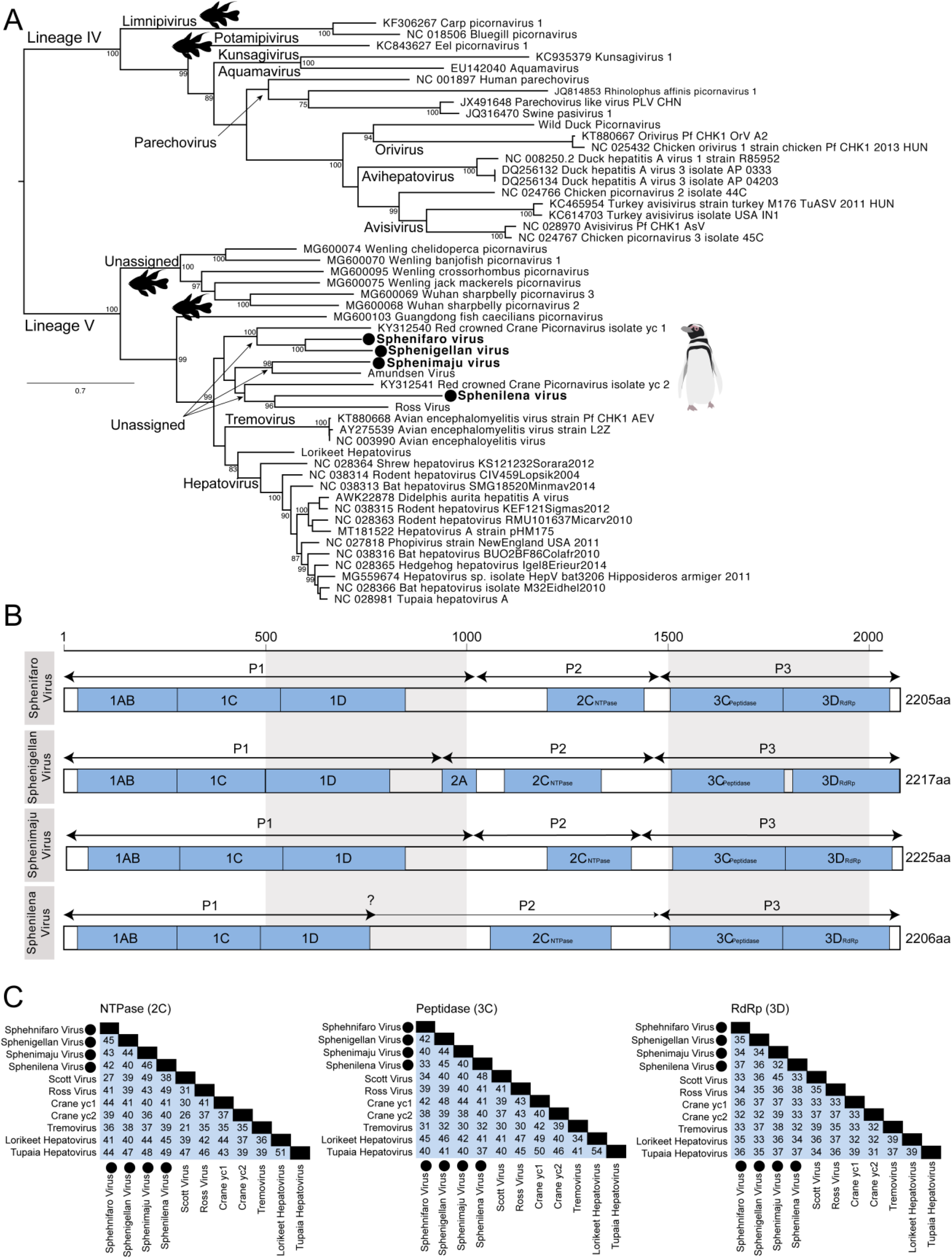
Genomic features of the novel picornaviruses revealed in this study. (A) Phylogeny of the P1 mature peptide of select genera of the *Picornaviridae*. We included members of lineage VI and V, as defined by (32), and the tree was rooted based on this lineage divergence. In addition to avian genera, we also included members of the *Limnipivirus* and *Potamipivirus* known to infect fishes and *Parechoviruses* known to include mammals. Viruses described here are adjacent to a filled circle. Genera and clades dominated by fish viruses have been indicated by a fish silhouette downloaded from phylopic.com and distributed by a Creative Commons attribution. Penguin silhouette was developed by M. Wille. Bootstrap values >70% are shown. The scale bar indicates the number of amino acid substitutions per site. A tree including all Picornavirus genera found in birds as is presented in Figure S1 (B) Genome arrangement for the four novel picornaviruses described in this study. Domains we were able to identify are indicated by filled box and place according to the amino acid position. Domains were identified using InterProScan. Detailed results of domain searches are presented in Table S3-S6. (B) Amino acid identity (percentage similarity) of the viruses revealed in this study and select reference sequences of the domains 2C_NTPase_, 3C_Peptidase_ and 3D_RdRp_

Phylogenetic analysis reveals that the most closely related fish picornaviruses fall into a sister clade to the avian and mammalian viruses of the *Hepatovirus*, *Tremovirus* and the recently described picornaviruses from Antarctic penguins, red-crowned cranes and the new viruses described here (Fig 1). Further, the viruses described in this study have less than 36% (22-36%) amino acid similarity to picornaviruses from fish across the P1. As such, the evidence suggests that the viruses revealed here are most likely *bona fide* avian viruses and do not derive from the prey.

To investigate the frequency of these four new viruses in other samples sequenced by high throughput sequencing, all libraries were mapped against each of these new viral genomes. In the sample containing Sphenimaju virus (is_034_015), there were over 5,000 reads mapping to the Sphenigellan virus. In the sample with both Sphenifaro virus and Sphenilena virus (is_034_016), there were over 1,000 reads mapping the Sphenigellan virus and Sphenimaju virus. Overall, there were 4 libraries with no evidence of any of the novel viruses described here, 2 with reads against only one of the novel viruses, and 6 libraries with evidence of more than one of these viruses (Table S2).

Using InterProScan domain prediction and our knowledge of *Picornaviridae* genomic features, we could predict the 3 main regions P1, P2, P3 for three of the viruses (Figure 1B). For Sphenilena virus, it was not possible to predict the exact P2 region location. Mature peptides 1AB, 1C and 1D could also be located, as well as 2C, 3C and 3D peptides for all 4 viruses. On the contrary, peptides 2A, 2B, 3A and 3B could not be predicted from the 4 polyproteins, however, most of the viral functional domains could be identified (Table S3 – S6): the viral coat, the helicase, the NTPase and the RNA dependant RNA-polymerase. The 4 assembled viral genomes and their annotations have been deposited to ENA, accession numbers: ERZ1667848, ERZ1667849, ERZ1667850, ERZ1667851.

### The four new picornaviruses are common among Magellanic penguins

To reveal the prevalence of the new viruses identified in this study, 107 faecal samples were screened by PCR for these 4 viruses. The samples were from 72 individual Magellanic Penguins, one Rockhopper penguin (*Eudyptes chrysocome chrysocome*), and three Kelp gulls (*Larus dominicanus*). Overall, at least one of these 4 viruses was detected in 28 samples (25%; 95% confidence interval [CI] 18-34%). All positive samples were from Magallanic Penguins, of which 22 were positive for at least one of the 4 viruses (21%, 95 CI 14-30%). The prevalence was highest for Sphenilena virus with 11 PCR positive samples. Sphenifaro virus, Sphenigellan virus and Sphenimaju virus were found in in 7, 7, and 9 Magellanic Penguin individuals, respectively. There was no statistically significant difference in prevalence between the viruses (*X*^2^ =3.01, df=3, p=0.3886) (Fig 3). Interestingly, samples from 7 individuals contained more than one of these viruses. Two individuals were positive for Sphenifaro virus and Sphenigellan virus, one individual for Sphenifaro virus and Sphenimaju virus, three individuals were positive for Sphenimaju virus and Sphenilena virus, finally, one sample contained 3 viruses: Sphenifaro, Spehnigellan, and Sphenilena virus.

Sequencing the ~460bp PCR product demonstrated sequence variability for each of the new viruses. There was an average pairwise identity of 91.6%, 96.5%, 98% and 91% for Sphenifaro, Sphenigellan, Sphenimaju, and Sphenilena virus, respectively. The pairwise similarity for Sphenimaju was higher (*i.e.* less diversity) than that for the other species, despite having a comparable number of PCR products (n=9) (Fig 4)

**Figure 2:**
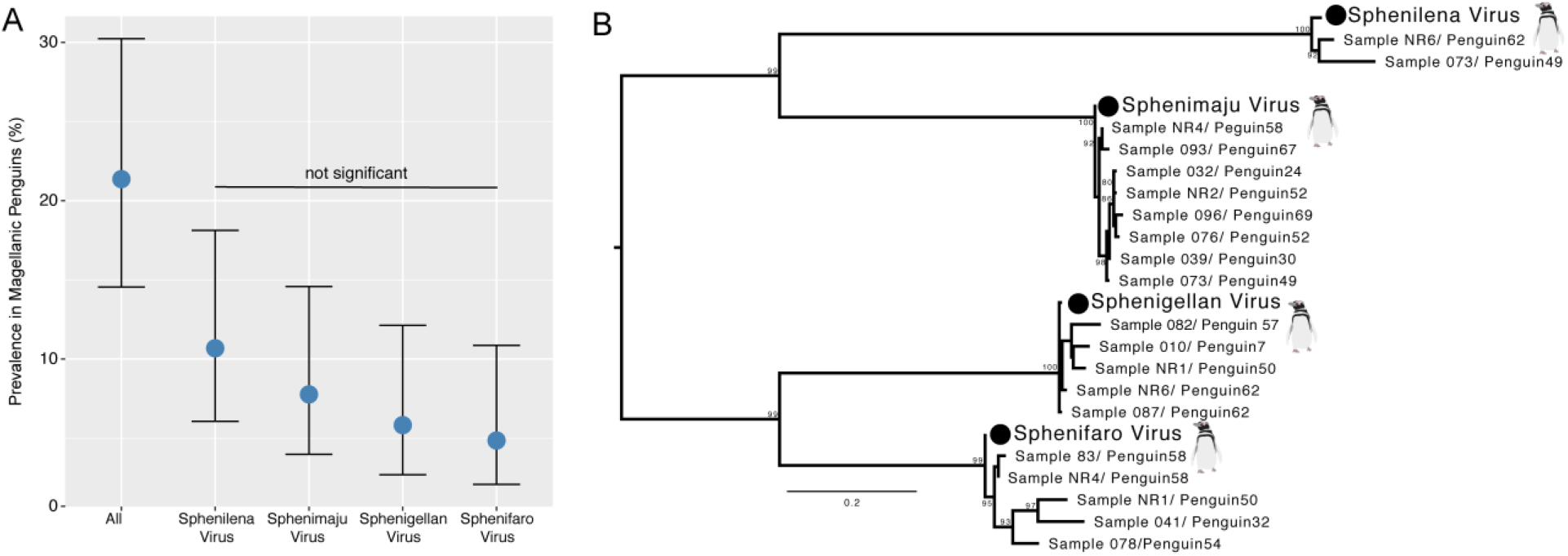
Diversity of four novel picornaviruses revealed through PCR. (A) Prevalence of each novel virus revealed in this study across samples collected from Magellanic Penguin individuals. The point estimate is presented as a filled circle and error bars correspond to 95% confidence intervals (B) Maximum likelihood tree of ~400bp PCR products sequenced from 20 positive samples. The tree was midpoint rooted for clarity only. Scale bar represents number of nucleotide substitutions per site. Viruses sequences assembled following metagenomic sequencing are indicated with a filled circle.

## Discussion

In this study we aimed to reveal the viral diversity of Magellanic Penguins sampled in Chile. By using high throughput sequencing we found four novel viruses belonging to the family *Picornaviridae*. These viruses are highly divergent and share less than 40% of the amino acid sequence in the P1 region with members of *Hepatovirus* and *Tremovirus* that they most closely resemble. The vast majority of the viral reads from most libraries mapped to these novel viruses indicate that these viruses were the dominant species in the faecal virome of the penguins. Further, through PCR screening we showed that over 20% of the sampled penguins in this study were shedding at least one of these viruses, and many individuals were co-infected with two or three of these new viruses. The viruses revealed here, in addition to novel and unassigned viruses from Antarctic Penguins (14) and Red-crowned Cranes (29) likely constitute at least three novel genera. This finding of very divergent avian picornaviruses forming one branch together with members of the genera Tremovirus and Hepatovirus shows on the need to further classify the genera with the *Picornaviridae* family into subfamilies or lineages, and illuminates that there are many more picornaviruses to be identified, especially from this part of the *Picornaviridae* phylogeny.

The charismatic Sphenisciformes have long been a target for virus surveillance, with early studies initiated in Antarctic Penguins in the 1970’s (9, 33). These studies were of course limited to screening for only described viruses, which until recently were viruses of poultry such as influenza A virus, Newcastle Disease virus, infectious bursal disease virus (*e.g.* (9, 33–36). As a consequence of this focus on poultry relevant virus surveillance, until very recently, few viruses had been described in penguin species, globally. While our viral catalogues are expanding, without detailed virus ecology studies, it remains unclear the role that penguins may play as hosts to an array of viruses. This is particularly true for viruses detected in penguins in rehabilitation centres (8). The high prevalence of the four novel viruses identified in this study, and the high frequency of co-infections identified, indicates that these penguins are likely an important reservoir hosts for these viruses. To understand the host range of these viruses, we would strongly encourage for the sampling and surveillance of not only other populations of Magellanic Penguins, but also other penguin species found in South America, and globally. Evidence for connectivity of penguin species and populations as hosts for viruses is sparse, however studies from avian avulavirus provide some clues. First, we have now seen repeated detection of Avian avulavirus 17, 18, 19 in three Antarctic Penguin species and across a number of colonies along the Antarctic peninsula. More interesting was the detection of avian avulavirus 10, first in Rockhopper Penguins from the Falkland/Malvinas Island (15), and more recently in Antarctic Penguins (14). This data suggests that there is capacity of viruses to be shared across multiple penguin species and locations. How these viruses may fit into the larger migratory flyways used by other birds in South America, such as Red Knot (*Calidris canutus*), is very unclear (37, 38). In addition to ecological questions, of importance would be to understand the route of transmission of these viruses. Given the detection in faeces, and the fact that faecal-oral route of transmission is very common in RNA viruses of wild birds, such as influenza A virus and avian avaulaviruses, we may speculate that these viruses are transmitted by the faecal oral route. This, however, would need to be confirmed by dedicated studies. Taken together, we would argue for, not only more sampling and metagenomic studies in these hosts, but also more consistent and repeated sampling to reveal both virus diversity and virus ecology.

Metagenomic tools have rapidly allowed for the expansion of described avian picornaviruses, but also redefined our understanding of the impact that these viruses have on their hosts. Prior to 2010, almost all described avian picornaviruses caused disease in their hosts. This includes duck hepatitis virus (genus *Avihepatovirus*) (39), turkey hepatitis virus (genus *Megrivirus*) (40) and avian sapelovirus (genus *Sapelovirus*) (41) which cause hepatitis in domestic ducks and turkeys, avian encephalomyelitis virus (genus *Tremovirus*) causing a neural disorder encephalomyelitis (42) in domestic gallinaceous birds, and a number of viral genera causing gastroenteritis in domestic galliforms (31). Indeed, a recent metagenomic study revealed that a chicken with gastroenteritis was shedding 6 picornaviruses, simultaneously, although it is unclear which, if any, of these viruses was causing the disease (31). There is no doubt that the viruses isolated from poultry cause disease, however metagenomic studies have now revealed more than 20 novel picornavirus species in wild birds, and on all occasions the sampled birds had no signs of disease (10, 13, 24–29), with the exception of Poecevirus causing avian keratin disorder (43). In this study, we demonstrated not only a high viral load (large proportion of reads), but also that a number of penguins were co-infected with at least two of the new picornaviruses described here without any obvious signs of disease. Similarly, in a metagenomic study, Wille *et al.* (2018) found Red-necked Avocets (*Recurvirostra novaehollandiae*) to be co-infected with nine different viruses, including 3 highly divergent picornaviruses. This suggests that wild birds are able to tolerate high viral loads and diversity in the absence of overt disease (44, 45). One hypothesis for the discordance in disease signs in wild birds as compared to poultry is that wild birds have a long history of host-pathogen co-evolution (24, 46). Mass produced chickens and ducks, which suffer from disease when infected with an array of virus species, are a relatively new host niche in evolutionary time (47, 48).

With more viral metagenomic studies being undertaken in under sampled hosts, such as fishes, many viral families once thought to exclusively infected birds and mammals now include members infecting fish and amphibians (49). Indeed, a number of picornaviruses have now been described in fish, and these viruses belong to genera falling within lineages previously dominated by viruses of mammmals and birds. The *Limnipvirus* and *Potamipivirus* genera are a case in point; these viruses fall within lineage IV (as defined by (32)) which includes a number of mammalian and avian infecting viral genera such as *Avihepatovirus* and *Parechovirus* (49–51). Shi *et al*. further described a number of fish viruses falling sister to established lineage IV viruses (*Hepatovirus* and *Tremovirus*) (49). Given that the viruses we characterized in this study were all identified in birds and are not in clades comprising fish viruses strongly suggests that we have detected avian viruses, rather than viruses of the diet of penguins. This distinction is especially important to clarify in samples collected from cloacae or faeces. These viruses of fish further demonstrate important evolutionary patterns in the lineage V viruses. That the phylogeny of the *Hepatovirus*, *Tremovirus*, and unassigned avian viruses and fish viruses generally follows the phylogeny of the hosts from which they are sampled suggests that there has been a process of virus-host co-divergence, that likely extends hundreds of millions of years. It further suggests that we may expect to find numerous avian picornaviruses that fall sister to the *Hepatovirus* and *Tremovirus* genera. It has been estimated that over 99% of viruses are still to be described (52), and therefore we anticipate a large number of picornaviruses are yet to be identified particularly from the incredible diversity of wild birds.

## Material and Methods

### Ethics Statement

The project was ethically approved by the Chilean National Forestry Corporation (CONAF) for the region of Magallanes, Chile (RESOLUCIÓN No: 517/2015).

### Study site

Sampling was carried out on the Magellanic penguin colony situated on the Magdalena Island in the Strait of Magellan in Chilean Patagonia (−52°55′10″S 70°34′34″W), from 19 - 21 November 2015. The colony comprises approximately 63,000 breeding pairs of penguins as the last count in 2007 (53).

Freshly deposited faecal samples (n=107) were collected from birds comprising 72 Magellanic penguins, 1 Southern Rockhopper penguin and, 3 Kelp Gulls. Some penguin faeces (n=30) were sampled on more than once. An additional 2 samples were collected from the soil around the colony. All samples were collected using sterile plastic tools and placed in 1 ml RNAlater (ThermoFisher) and stored at room temperature for up to 72h prior to storage in −80 °C. For eight penguins, a duplicate sample was stored dry in sterile tubes without RNAlater for approximately 5h at 8 °C prior to storage in −80 °C. Samples remained at −80 °C until processing.

### Sample preparation

An initial 12 faecal samples from Magellanic penguins were prepared for viral metagenomic sequencing. Eight of the samples had been stored in RNAlater (ID:s 1,11,34,36,51,81,87,88) and 4 samples were stored in dry tubes without RNAlater (NR2, NR4, NR5, NR6). Dry samples were treated with 20 U RNase I (Epicentre) prior to RNA extraction.

All samples were homogenized in 800μl PBS using OmniTip Homogenizer (Omni International Inc) for 15 sec and then placed on ice for 30 sec followed by 15 sec homogenization. The homogenate was centrifuged at 4,000 rpm for 10 minutes, thereafter the supernatant was transferred to a 0.65μm filter (Millipore) and centrifuged at 12,200 rpm for 4 minutes. The filtrate was further transferred to a Ultrafree-CL GV 0.22 μm Sterile filter column and centrifuged at 4,800 rpm for 4 minutes. Thereafter RNA was extracted using Trizol and choloroform and the RNA was cleaned up using RNAeasy (Qiagen) according to protocol. Qubit HS RNA assay was used for quantification.

The extracted RNA was amplified using Ovation RNA-Seq v2 (NuGEN) according to the manufacturer’s recommendations. The final product was purified with GeneJET PCR purification kit (ThermoFisher) followed by Qubit HS DNA assay for quantification.

A negative PBS control samples was included throughout the laboratory workflow.

### Library preparation and sequencing

Sequencing libraries were constructed using the AB Library Builder System (Ion Xpress™ Plus and Ion Plus Library Preparation for the AB Library Builder™ System protocol, ThermoFisher) and size selected on the Blue PippinTM (Sage Science). Library size and concentration were assessed by a Bioanalyzer High Sensitivity Chip (Agilent Technologies) and by the Fragment Analyzer system (Advanced Analytical). Template preparation was performed on the Ion Chef™ System using the Ion 520 & Ion 530 Kit-Chef (ThermoFisher). Samples were sequenced on 530 chips using the Ion S5™ XL System (ThermoFisher).

### Bioinformatics analysis

The obtained reads were trimmed by quality in 5’ and 3’ and filtered by mean quality using PRINSEQ (v 0.20.4) with a PHRED quality score of 20 (54). The good quality reads were used to produce *de-novo* assemblies with Megahit version 1.1.1 (55). A taxonomic classification of the obtained contigs was performed by running Diamond (v 0.9.6) (56) against the non-redundant protein database (release April 2019) and using the LCA algorithm from Megan 6 (v6.11.7) (57) to visualise the classification of each contig. A taxonomic classification at the read level was also performed using Diamond (blastx + LCA, using the output format 102). The classified output table was converted into a Kraken report to allow visualisation with the R package Pavian (58).

### Comparative genomics and phylogenetic

We interrogated the 4 contigs representing near full length picornavirus genomes (>6000bp). Gene prediction was performed using ORFfinder (https://www.ncbi.nlm.nih.gov/orffinder/) and protein domains prediction using InterProScan (59). Reads were subsequently mapped back to viral contigs using the Burrows Wheeler Aligner BWA-MEM (60).

Amino acid sequences of the polyprotein were aligned using MAFFT with the E-INS-I algorithm (61). Reference genomes for the *Picornaviridae* tree were limited to those genera containing avian (and mammalian) genera as per Wille *et al*. 2019b. Gaps and ambiguously aligned regions were stripped using trimAL (62). The most appropriate amino acid substitution model was then determined for each data set, and maximum likelihood trees were estimated using IQ-TREE (63).

### Annotation and file conversion for submission of annotated viral genome to ENA

For generating EMBL flat files to submit the annotated viral genome to ENA, we have used several tools. First, we used Prokka (64), for generating a GFF file with the polyprotein coding sequence (CDS) with some manual modification if required.

Using scripts from AGAT (Another Gff Analysis Toolkit; https://github.com/NBISweden/AGAT, version v0.5.0), the corrected GFF files were standardised with ‘agat_convert_sp_gxf2gxf.pl’ and proteins were extracted using ‘agat_sp_extract_sequences.pl’. The extracted polyproteins were then run through InterProScan online and TSV files containing the domain annotations were downloaded. The InterProScan annotation could be integrated into the GFF file using ‘agat_sp_manage_functional_annotation.pl’. In order to submit the annotated viral genomes to ENA, the GFF files were converted into EMBL file using EMBLmyGFF3 (65).

### Ascertaining the prevalence of four novel picornaviruses

RNA was extracted from the 107 faecal samples using QIAamp Fast DNA Stool mini kit (Qiagen), followed by cDNA synthesis using the High-Capacity cDNA Reverse Transcription (Applied Biosystems). Custom primers were designed for each of the four hepato-like viruses revealed (Table S7). For Sphenifaro and Sphenigellan viruses a nested PCR approach was employed, and for Sphenimaju and Sphenilena viruses a semi-nested approach was employed. The 50 μl PCR reaction mix contained 1xTaq Buffer, 2.25mM of MgCl_2_ (Applied Biosystems), 0.2mM of dNTP (Roche), 1 U of Taq Polymerase (Roche), 1.5mM of each primer and 5μl of cDNA. The same reaction mix was used in the nested PCRs with 5μl of the first PCR product as template. The PCR reactions were run with primary denaturation at 94°C for 4 minutes followed by 40 cycles of denaturation at 94°C for 20 seconds, annealing at 54°C for 30 seconds followed by polymerization at 72°C for 90 seconds. The PCR products were visualized on 1.5% agarose gel electrophoresis and amplified products were purified using QIAquick PCR purification kit (Qiagen) according to manufacturer’s protocol. Purified amplicons were sent to Eurofins Genomics (Germany) for Sanger sequencing.

Viral prevalence was calculated using the *bioconf()* package and statistically evaluated using a Chi squared test in R v 3.5.3 integrated into RStudio 1.1.463.

## Acknowledgments

Financial support was obtained from the DEANN project, Marie Curie Action funded by the European Commission – Grant agreement ID: 612583 and The Swedish Research Council for Environment, Agricultural Sciences and Spatial Planning, Formas (grant number 221-2012-586).

We acknowledge personnel at the Chilean National Forestry Corporation (CONAF) and assistance by the National Genomic Infrastructure (NGI), which is a part of the Science for Life Laboratory (SciLifeLab), as well as the Swedish National Infrastructure for large Scale Sequencing (SNISS) initiatives. NGI is financially supported by SciLifeLab, Knut and Alice Wallenberg foundation, and the Swedish Research Council.

We acknowledge the SLU Bioinformatics Infrastructure (SLUBI) for providing computing resources and supporting JH.

MW is supported by an Australian Research Council DECRA Fellowship (DE200100977).

We acknowledge the support with molecular methods provided by Alexandra Corduneanu at University of Agricultural Sciences and Veterinary Medicine in Romania and Fernando Pinto Swedish at University of Agricultural Sciences in Sweden.

